# Lipin-1 contributes to IL-4 mediated macrophage polarization

**DOI:** 10.1101/2019.12.23.887109

**Authors:** Sunitha Chandran, Robert M. Schilke, Cassidy M.R. Blackburn, Aila Yurochko, Rusella Mirza, Rona S. Scott, Brian N. Finck, Matthew D. Woolard

## Abstract

Macrophage responses contribute to a diverse array of pathologies ranging from infectious disease to sterile inflammation. Polarization of macrophages determines their cellular function within biological processes. Lipin-1 is a phosphatidic acid phosphatase in which its enzymatic activity contributes to macrophage pro-inflammatory responses. Lipin-1 also possesses transcriptional co-regulator activity and whether this activity is required for macrophage polarization is unknown. Using mice that lack only lipin-1 enzymatic activity or both enzymatic and transcriptional coregulator activities from myeloid cells, we investigated the contribution of lipin-1 transcriptional co-regulator function towards macrophage wound healing polarization. Macrophages lacking both lipin-1 activities did elicit IL-4 mediated gene expression to levels seen in either wild-type or lipin-1 enzymatically deficient macrophages. Furthermore, we provide evidence that the lack of myeloid-associated lipin-1 transcriptional co-regulator activity leads to impaired full thickness excisional wound healing. Our study demonstrates that lipin-1 transcriptional co-regulatory activity contributes to macrophage polarization and the macrophage contribution to wound healing *in vivo*.

## INTRODUCTION

Macrophages are innate immune cells that mediate tissue homeostasis by polarizing into unique phenotypes that range from pro-inflammatory to wound healing. Macrophage cellular responses restore normal tissue function. Defects in macrophage polarization can influence numerous disease pathologies including infectious disease, atherosclerosis, tumor growth, and impaired wound closure. Activation of macrophages via pattern recognition receptors (i.e Toll like receptors) or through pro-inflammatory cytokine receptors (i.e. IFN-γ- or TNF-α receptor) lead to pro-inflammatory activities [1]. Conversely, IL-4, IL-10, IL-13 or TGF-β stimulation of macrophages promotes wound healing activities The phosphorylation/activation of STAT-6 and subsequent activation of peroxisome proliferator-activated receptor γ (PPARγ) is crucial to macrophage wound-healing polarization [2]. PPARγ activation via ligand binding and association with co-activators leads to both trans-repressive and transactivating activity [3]. In macrophages, PPARγ transrepresses NF-κB and STAT-1 at the promoters of pro-inflammatory cytokines such as TNF-α [4], and promotes the expression of genes associated with lipid metabolism [2], driving macrophage wound-healing polarization [5].

Lipin-1 belongs to the evolutionarily conserved three-member lipin family (lipin-1, −2, and −3) in mammals. Lipins enzymatically convert phosphatidate into diacylglycerol via dephosphorylation and among lipin family proteins, lipin-1 exhibits the highest phosphatidate-specific phosphohydrolase activity [6]. We and others have shown that expression of a hypomorphic lipin-1 protein that lacks enzymatic activity attenuates pro-inflammatory macrophage responses by regulating glycerolipid synthesis [7; 8; 9]. Lipin-1 enzymatic activity within macrophages contributes to disease pathogenesis of atherosclerosis, colitis, colon cancer, and LPS induced inflammation (reviewed in [10]). The overarching mechanism is likely due to lipin-1-mediated diacylglycerol production leading to protein kinase C and AP-1 transcription factor activation driving pro-inflammatory macrophage activities [7; 8; 9]. In addition to acting as a lipid phosphatase, lipin-1 also independently acts as a transcriptional co-regulator by interacting with various DNA-bound transcription factors. Lipin-1 augments PPAR activity to promote adipogenesis in adipocytes and promotes beta-oxidation and while suppressing very low-density lipoprotein production in hepatocytes [11; 12; 13; 14; 15]. It is unknown if lipin-1 transcriptional co-regulator activity is involved in regulating wound healing activity in macrophages. Our data provides evidence that lipin-1 transcriptional co-regulator activity contributes to IL-4 mediated macrophage wound healing function.

## MATERIAL METHODS

### Animals

All animal studies were approved by the LSU Health Sciences Center-Shreveport institutional animal care and use committee. All animals were cared for according to the National Institute of Health guidelines for the care and use of laboratory animals. After wounding, all mice were housed in individual filter-topped sterile cages, provided with sterile water and food *ad libitum*.

All animals used in this study were 8 to 10 weeks old mice. Mice lacking lipin-1 enzymatic activity from myeloid cells (lipin-1^mEnzy^KO) were generated as previously reported [9]. Briefly, mice with exons 3 and 4 of the *Lpin1* gene flanked by LoxP sites (genetic background: C57BL/6J and SV129; generously provided by Brian Finck and Roman Chrast) were crossed with C57BL/6J *LysM*-Cre transgenic mice purchased from Jackson Laboratory (Bar Harbor, ME). Exon 3 encodes the translational start site of lipin 1; however, deletion of this exon led to enforcement of an alternative start site causing expression of a truncated lipin 1 protein lacking 115 amino acids [16]. The truncated protein lacks phosphatidic acid phosphohydrolase activity but retains transcriptional regulatory function. Mice fully lacking lipin-1 from myeloid cells (lipin-1^m^KO) were generated by crossing mice with exon 7 of the *Lpin1* gene flanked by LoxP sites (genetic background: C57BL/6J and SV129; generously provided by Brian Finck [17]) with C57BL/6J *LysM*-Cre transgenic mice purchased from Jackson Laboratory (Bar Harbor, ME). Deletion of exon 7 leads to frameshift, premature stop codon insertion, and a complete loss of lipin 1 protein [17]. Age matched lipin-1^flox/flox^ littermate mice were used as controls.

### Excisional wound healing model

Mice were anesthetized by 3% isoflurane (NDC, 14043-704-06) and clippers were used to remove hair from the dorsum. Exposed skin was disinfected with chlorohexidine swabs. Dorsal skin was folded, raised cranially, and mice were laterally positioned. Symmetric full thickness wounds were created using a sterile 5mm biopsy punch (Integra) [18]. Gross images were taken and percentage of wound closure was assessed using a digital caliper at 0, 2, 5, 7, 9, 12, and 14-days post-surgery and expressed as [(area of original wound – area of actual wound)/area of original wound]x100. After initial wounding, analgesic cream was applied to wounds (Aspercreme, Cattem, 0078940). Mice were routinely monitored for weight loss or any other type of distress until the end of the study.

### Generation of bone marrow-derived macrophages

Bone marrow-derived macrophages (BMDMs) were generated from lipin-1^mEnzy^KO, lipin-1^m^KO and littermate control mice as previously described [19]. Briefly, femurs were excised under sterile conditions and flushed with Dulbecco’s modified Eagle’s Knock out medium (DMEM; Gibco, 10829) supplemented with 5 % fetal bovine serum (Atlanta biologicals, S11150), 2mM L-glutamax (Gibco -35050-061), 100 U/ml penicillin-streptomycin (Cell gro, 30-604-CI), 1mM sodium pyruvate (Cell gro, 25-060-CI), and 0.2 % sodium bicarbonate (Quality biological, 118-085-721). Red blood cells were lysed using ammonium chloride-potassium carbonate (0.15 M NH4Cl, 10 mM KHCO3, 0.1 mM NA2EDTA, adjusted to pH 7.2 and filter sterilized in 0.22 μm filter) lysis (ACK) followed by PBS wash. Isolated cells were incubated in sterile petri dishes for 7 to 10 days in BMDM differentiation medium - DMEM (Gibco, 11965-092) supplemented with 30% L-cell conditioned medium, 20% fetal bovine serum (Atlanta biologicals, S11150) at 37°C and 5% CO_2_. Once cells were 80% confluent, they were collected using 10 mM EDTA, pH 7.6 treatment. 10^6^ cells were seeded for RNA extraction and 5×10^5^ cells for protein isolation and flow cytometry analysis. After 4 hours of seeding, cells were treated with 0 or 20 ng/ml IL-4 (R&D Biosystems, 404-ML-050) for various times.

### Flow cytometry

#### Interleukin-4 Receptor Staining

BMDMs were incubated with CD16/CD32 (e-Bioscience, 14-0161-86) for 20 min. Cultured cells were incubated with PECy7 conjugated CD11b (e-Biosciences, 25-0112-81, clone M1/70), and PE conjugated IL-4R (Biolegend, 144803) for 30 minutes in the dark. Cells were then fixed with 4% formaldehyde and analyzed using BD LSRII (San Jose, CA).

#### Immune Composition Staining

Spleens were homogenized in FACS wash buffer (1% bovine serum albumin, 1 mM EDTA, and 0.1% sodium azide in phosphate buffered saline). The spleens were strained with a 40 μm cell strainer (Falcon, 352340) followed by centrifugation at 300Xg for 5 minutes. The supernatant was decanted and splenocytes were dislodged in 3 mLs of ACK lysis buffer. Splenocytes were incubated on ice for 5 minutes. Splenocytes were washed in FACS wash buffer then centrifuged. The pellet was re-suspended in FACS wash buffer, strained (Falcon, 352340), and counted. Splenocytes were adjusted to 1×10^6^ cells/mL in RPMI. Blood was collected in EDTA coated tubes. 100 ul of blood was lysed in 3 mls of ACK lysis buffer and then washed with FACS wash buffer. The entire sample of blood cells were stained. Splenocytes and blood cells were incubated with CD16/CD32 (e-Bioscience, 14-0161-86) for 20 min. After blocking, cells were stained with a cocktail of antibodies: AF700 conjugated CD45.2 (Biolegend,109821,clone104), BV605 conjugated CD3 (Biolegend,100237,clone17A2), BV786 conjugated CD11c (BD Biosciences,563735,cloneHL3), PECy7 conjugated CD11b (eBiosciences, 25-0112-81, clone M1/70), PEe610 conjugated CD19 (eBiosciences,61-0193-80,clone eBio1D3), FITC conjugated Ly6G (BD Biosciences,551460,clone1A8), PE conjugated Ly6C (eBiosciences,12-5932-80,clon eHK1.4) and APC-Cy7 conjugated CD115 (Biolegend,135532,cloneAFS 98). Appropriate F Minus One Controls were used to correct background and exclude spectral overlap staining. Compensation control (Comp Bead, Invitrogen, 01-2222-42) were used. Flow cytometry analysis was performed using BD LSRII (San Jose, CA). Data analysis was done using FCS express (Denovo Software) and NovoExpress (AceaBio).

#### Western blot

Cells were lysed in 1X NuPage LDS sample buffer (containing 100 mM dithiothreitol (DTT; Life Technologies), 1X protease inhibitor cocktail (Thermo Scientific), 1X phosphatase inhibitor cocktail 1 (Sigma Aldrich), and 1X phosphatase inhibitor cocktail 3 (Sigma Aldrich)). Protein concentration was determined by Peirce® 660 nm Protein Assay (Thermo Scientific) and 20 μg of each sample was separated using 4 to 12% polyacrylamide NuPAGE Novex gel (Invitrogen) run at 200V for 55 minutes. Semidry transfer (Novex, SD1000) was performed for 45 minutes at 20 V onto a polyvinylidene difluoride (Immobilon-FL) membrane (EMD Millipore). The membranes were further blocked for 1 hour at room temperature using 10% Li-Cor blocking buffer (Li-Cor Biosciences) and incubated overnight with primary antibodies for Lipin-1 (CST #14906), P-STAT6 (CST #56554), STAT6 (CST #5397), and GAPDH (CST #2118). Goat anti-rabbit HRP secondary antibody (Jackson #111-035-144) was added to the membranes and incubated for 2 hours at room temperature. Membranes were washed three times with tris buffered saline with tween and incubated in ImmunoCruz Western blotting luminol reagent (Santa Cruz, sc-2048) for 1 minute. Images were captured using an Amersham Imager 680 (GE Healthcare Bio-Sciences). Densitometry was performed using IQTL 8.1 (GE Healthcare Biosciences). Bands of interest were normalized to GAPDH.

#### Quantitative real time PCR

BMDMs were treated with IL-4 (R&D Biosystems, 404-ML-050) for 4 hours and mRNA was extracted from the cultured cells using RNeasy Mini Kit (Qiagen – 74106) as per manufacturer’s instructions. cDNA template was generated using qScript cDNA SuperMix (Quantabio, 95048). qRT-PCR was performed in a Biorad iCycler with SsoAdvanced Universal SYBER Green SuperMix (Biorad, 172-5271). Primers (Supplemental Table 1) were obtained from primer bank database. Primer specificity was confirmed using primer BLAST and by verifying the presence of a single peak in melt curve analysis. Results were expressed as fold change relative to IL-4 treated WT cells by 2^(−ΔΔCt)^ method after normalizing with GAPDH.

**Table 1.**
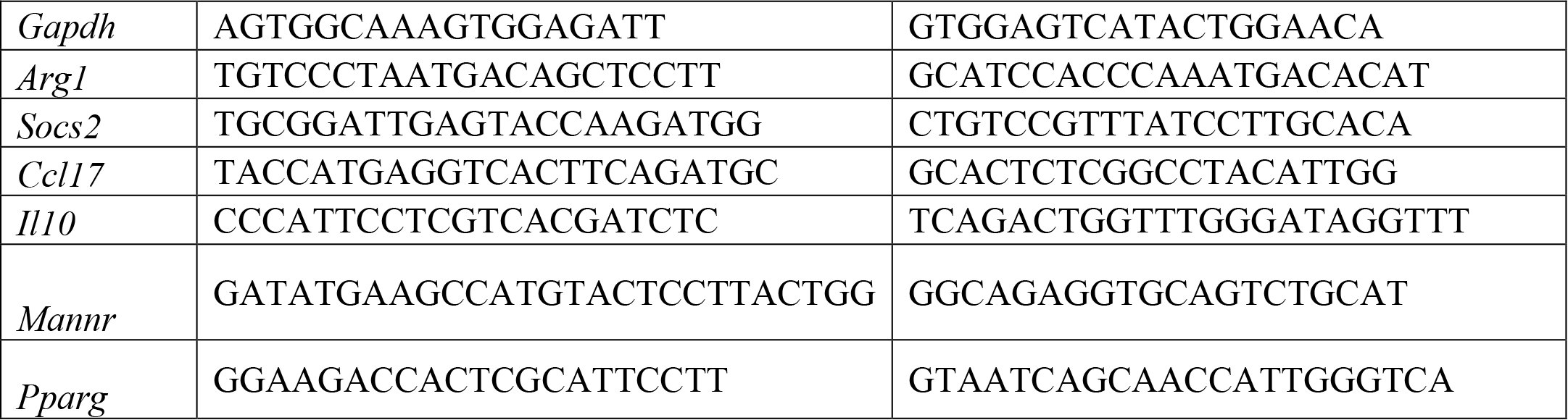

#### Phagocytosis assay

BMDMs (5×10^5^ cells) were cultured on sterile coverslips in culture wells and treated with 20 ng/ml IL4 for 24 hours. Culture medium was then replaced with DMEM, containing pHrodo™ green Zymosan A BioParticles® (Thermoscientific, P35365) such that each well receives 0.1mg zymosan particles. BMDMs were allowed to phagocytose for 1 hour under dark incubation and then the assay was stopped by cold PBS wash. Cell seeded on coverslips were then fixed using 4% formaldehyde. Cells on cover slips were washed three times and then stained with DAPI slowfade (Invitrogen, S36938). Immunofluorescent images were taken using Olympus BX51 and evaluated using Image J (1.50a) analysis software. Phagocytic efficiency for each image was calculated by dividing the total number of fluorescent beads by the total number of nuclei in the fluorescent image, thus giving average number of beads per cell. Experiment was performed 3 times with 4 random images per group (n=12).

#### Histology

Wound area was carefully excised at 2, 5 and 14 days after wounding and fixed in 10% neutral buffered formalin followed by paraffin embedding. 5 μm thick sections were cut from formalin-fixed paraffin-embedded tissue blocks. Sections were rehydrated, followed by hematoxylin-eosin (H&E) staining and dehydration. Stained sections were then imaged using Olympus BX51. 4x images were compared between each group to assess wound healing. Morphological score of inflammation: Evaluation of cellular infiltrate (polymorphonuclear and mononuclear cells) was done on H&E stained sections using the 10 x objectives. The cells were counted at the wound bed and scored as 0, 1, 2, 3 (absence of inflammation, Discrete-presence of few inflammatory cells, Moderate-many inflammatory cells and Severe-exaggerated inflammatory cellularity respectively) for whole skin. The cellularity of the overlying crust or scab was excluded from the score. The scab was made of fibrin and polymorphonuclear cells. The scab was interpreted as either thin (scored as 1) or thick (Scored as 2) based on their morphological appearance on H & E sections. Scoring was done blinded.

#### Cytometric Bead Array

Serum cytokine concentration was measured using Biolegend LEGENDplex (Biolegend Mouse inflammation Panel #740446). The assay was performed according to the manufacturer’s instructions, and all samples were run in duplicate. Data was analyzed using the LEGENDplex Data Analysis Software.

#### Statistical analysis

GraphPad Prism 5.0 (La Jolla, CA) was used for statistical analyses. Student T Test analysis was used for comparison between two data sets. All other statistical significance was determined using a one-way ANOVA analysis of variance with a Dunnett’s post-test, P ≤ 0.05

## RESULTS

### Lipin-1 contributes to IL-4 elicited gene expression

Pro-inflammatory response in macrophages is influenced by lipin-1, but if and how lipin-1 contributes to wound healing responses by macrophages is unknown. We have previously generated lipin-1^mEnzy^KO mice that express a truncated lipin 1 protein lacking lipin-1 enzymatic activity but retaining transcriptional regulatory function in myeloid cells [9]. Here, we generated lipin-1^m^KO that lack the entire lipin-1 protein in myeloid cells. Comparing results between lipin-1^mEnzy^KO mice and lipin-1^m^KO mice allows us to determine the contribution of lipin-1 enzymatic activity and infer the contribution of lipin-1 transcriptional coregulator activity on macrophage function. We have previously demonstrated the ability to generate BMDMs from lipin-1^mEnzy^KO mice and confirmed their phenotype [9]. We confirmed that the loss of lipin-1 did not inhibit BMDM generation based on CD11b staining by flow-cytometry (Fig 1A). Western Blot analysis of proteins collected from cultured BMDMs demonstrated roughly an 85% reduction of lipin-1 protein in lipin-1^m^KO BMDMs (Fig 1B). Having generated macrophages lacking lipin-1, we investigated the contribution of lipin-1 to IL-4 mediated gene expression. BMDMs from lipin-1^mEnzy^KO, lipin-1^m^KO, and littermate controls were stimulated with 20ng/ml of IL-4 for 4 hours. mRNA was isolated and analyzed for the expression of several wound-healing associated genes: *Arg1, Socs2, Ccl17, Mannr, Il10, and Pparg* [20]. We included littermate controls for both strains, however, no differences were noted between each littermate strain controls, therefore littermate controls were grouped together as wild type. IL-4 stimulation promoted the expression of several wound healing associated genes in wildtype, lipin-1^mEnzy^KO and lipin-1^m^KO BMDMs (Fig 1C). However, IL-4 elicited gene expression was significantly lower in lipin-1^m^KO BMDMs compared to either wild type or lipin-1^mEnzy^KO BMDMs. These results suggest that lipin-1 transcriptional co-regulatory activity influences IL-4-mediated gene expression by macrophages.

**Figure 1.**
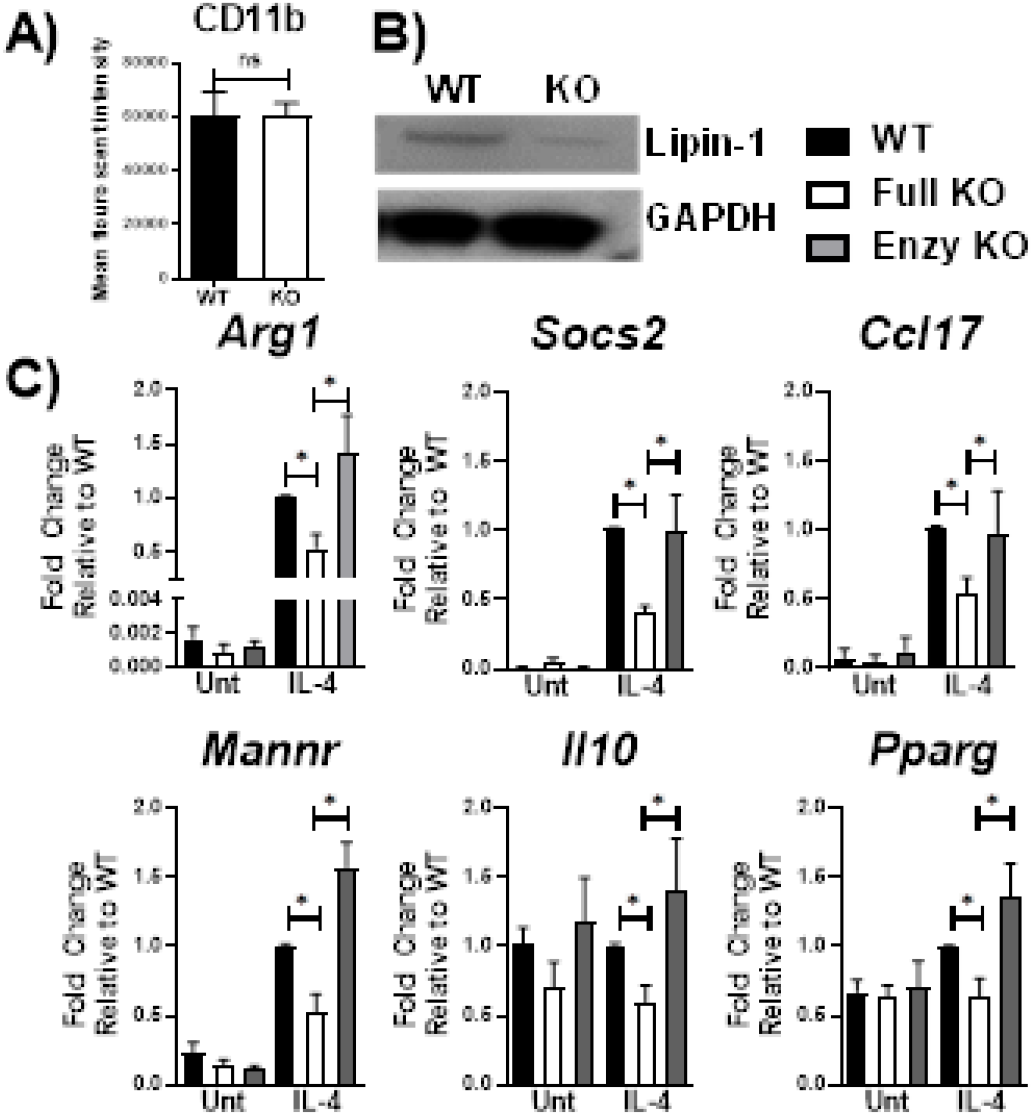
Lipin-1 promotes IL-4 mediated gene expression. A) Flow cytometry was used to quantify CD11b surface expression of BMDMs of lipin-1^m^KO and littermate controls. B) Lipin-1 was quantified by Western blot analysis representative image shown. C) BMDMs generated from lipin-1^m^KO, lipin-1^mEnzy^KO and their respective littermate control mice. BMDMs were stimulated with 20ng/ml IL-4 for 4 hours. mRNA was isolated and wound healing associated genes were quantified by qRT-PCR. No difference was noted between littermate controls as such they were combined in WT. (n≥3, *=p≤0.05)

### Lipin-1 does not regulate influence surface expression of IL-4 receptor expression or STAT6 phosphorylation

Lipid membrane composition can influence both the localization of receptors and/or signaling through those receptors [21]. Lipin-1 is a regulator of glycerol lipid synthesis and the loss of lipin-1 may cause loss of either IL-4 receptor surface expression or signaling through the IL-4 receptor, thus resulting in impaired responses to IL-4. Flow cytometric evaluation of the surface expression of IL-4 receptor showed no difference between wild type and lipin-1^m^KO BMDM (Fig 2A). Ligand binding of the IL-4 receptor-α (IL4Rα) triggers tyrosine phosphorylation at the cytoplasmic tail to facilitate recruitment and subsequent tyrosine phosphorylation of STAT6 by JAK1/JAK3 pathway [22]. Wildtype and lipin-1^m^KO BMDMs were stimulated with IL-4 for 30 minutes and protein was collected. Total STAT6 and phosphorylated STAT6 was measured by Western blot analysis. Similar levels of STAT-6 phosphorylation was observed between wild type and lipin-1^m^KO BMDMs (Fig 2B). These results indicate that defective wound healing polarization was likely not due to alteration in IL-4 binding to the IL-4 receptor and subsequent signaling.

**Figure 2.**
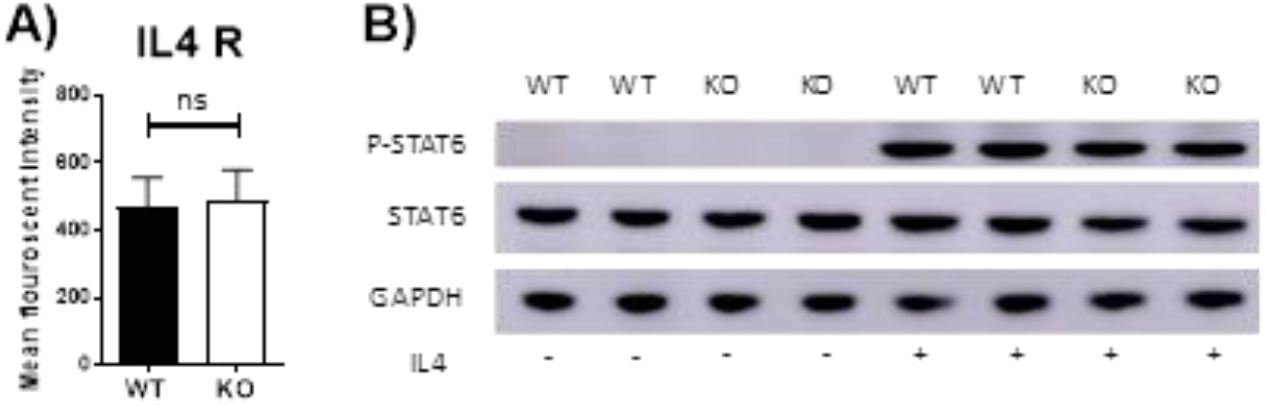
Lipin-1 does not regulate IL-4 mediated STAT6 phosphorylation. A) Flow cytometry was used to quantify surface expression of IL-4 in unstimulated BMDMs from Lipin-1mKO and litter mate controls n≥3. B) BMDMs from Lipin-1mKO and litter mate controls were stimulated with 20 ng/ml IL-4 for 30 minutes. Protein was isolated and p-STAT6, STAT6 was quantified by Western blot analysis. Representative blot shown. n≥3

### Lipin-1 is required for phagocytosis

The reduction in wound healing-associated genes in response to IL-4 suggests that lipin-1 contributes to macrophage wound healing function. Macrophages with a wound healing phenotype have increased phagocytic capabilities [23]. We investigated the ability of BMDMs to phagocytize zymosan beads. We either mock treated or IL-4 treated BMDMs from lipin-1^m^KO or litter mate control for 24 hours. We then fed the macrophages pHrodo™ green Zymosan A BioParticles for one hour. These particles do not fluoresce at 7.6 pH but do at acidic pH, making it easier to identify internal particles. We then imaged using fluorescent microscopy and quantified average number of particles per cell. IL-4 stimulated lipin-1^m^KO BMDMs had fewer particles per cell than wild type BMDMs (Fig 3). These results further implicate the importance of lipin-1 to macrophage wound healing function.

**Figure 3.**
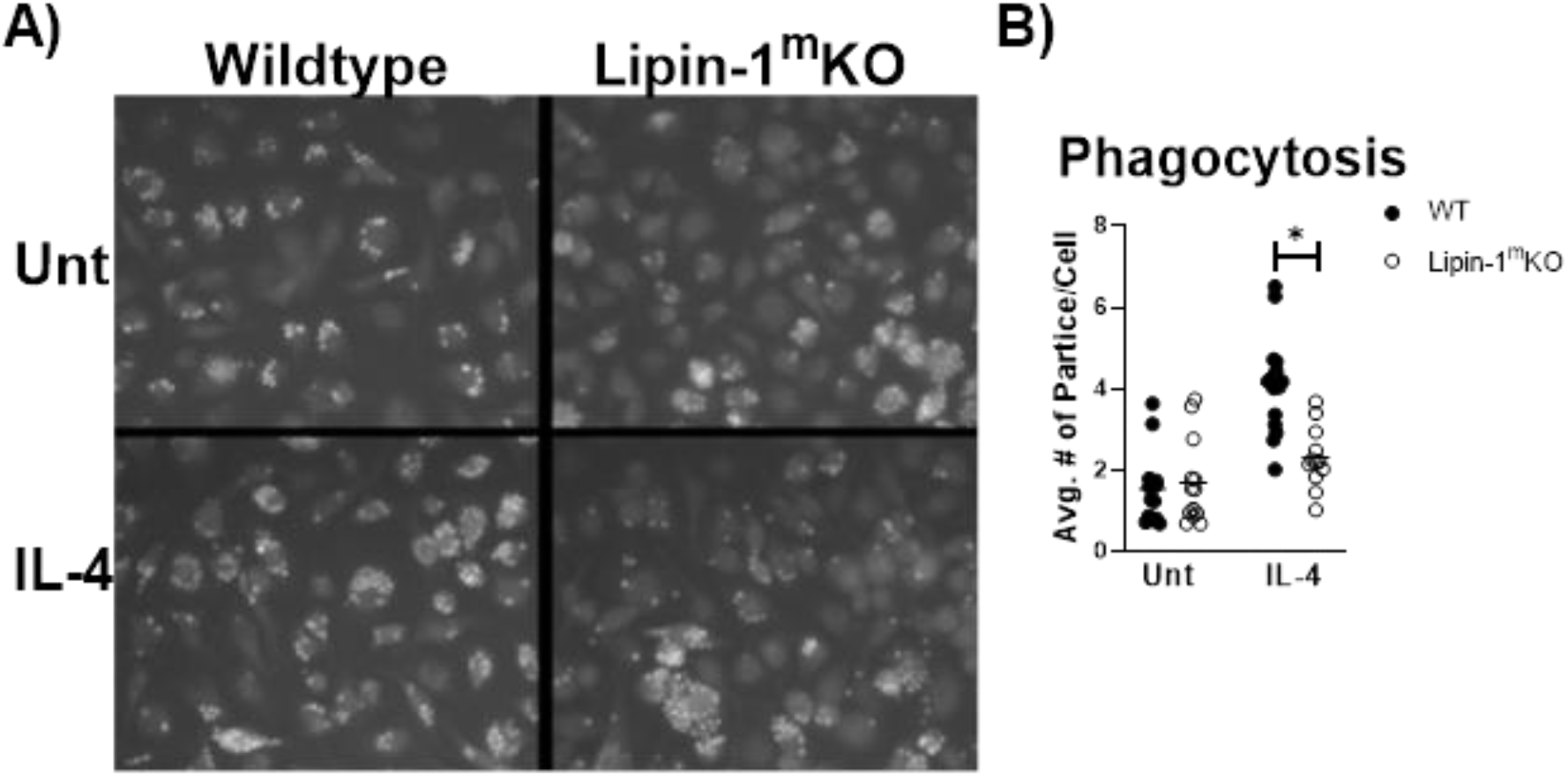
Lipin-1 contributes to IL-4 enhancement of phagocytosis. A) Representative microscopic images of BMDMS from lipin-1^m^KO and littermate control mice fed pHrodo-Green zymosan particles. B) Quantification of number of beads (zymosan beads) divided by number of nuclei in a given image. Experiment was performed 3 times with 4 random image panels taken per group total of 12 images. Each dot represents analysis of a single image. (n≥12, *=p≤0.05)

### Myeloid-associated lipin-1 contributes to wound healing *in vivo*

Our *in vitro* studies suggest that lipin-1 contributes to IL-4 mediated macrophage polarization. We next wanted to determine if these *in vitro* differences contribute to *in vivo* processes as well. Polarization of macrophage to a wound healing phenotype is required for proper wound closure in a full excision wounding model [24; 25]. We decided to investigate if the loss of myeloid-associated lipin-1 would alter wound closure. We performed full excision wounding on lipin-1^m^KO, lipin-1^mEnzy^KO and their respective littermate controls. We monitored wound closure at early (day 2 and day 5), middle (day 7 and day 9), and late (day 12 and day 14) stages of wound healing. Lipin-1^m^KO mice had initial delay in wound healing (days 2, 5, and 7) in lipin-1^m^KO as compared to litter mate controls (Fig 4A&B). 9 days after wounding, wounds were of comparable size between lipin-1^m^KO mice and litter mate controls. In contrast, lipin-1^mEnzy^KO did not differ from littermate controls in wound healing (Fig 4C&D) at any stage of healing. Thus, these results highlight the potential contribution of myeloid-associated lipin-1 transcriptional co-regulator activity in wound healing.

**Figure 4.**
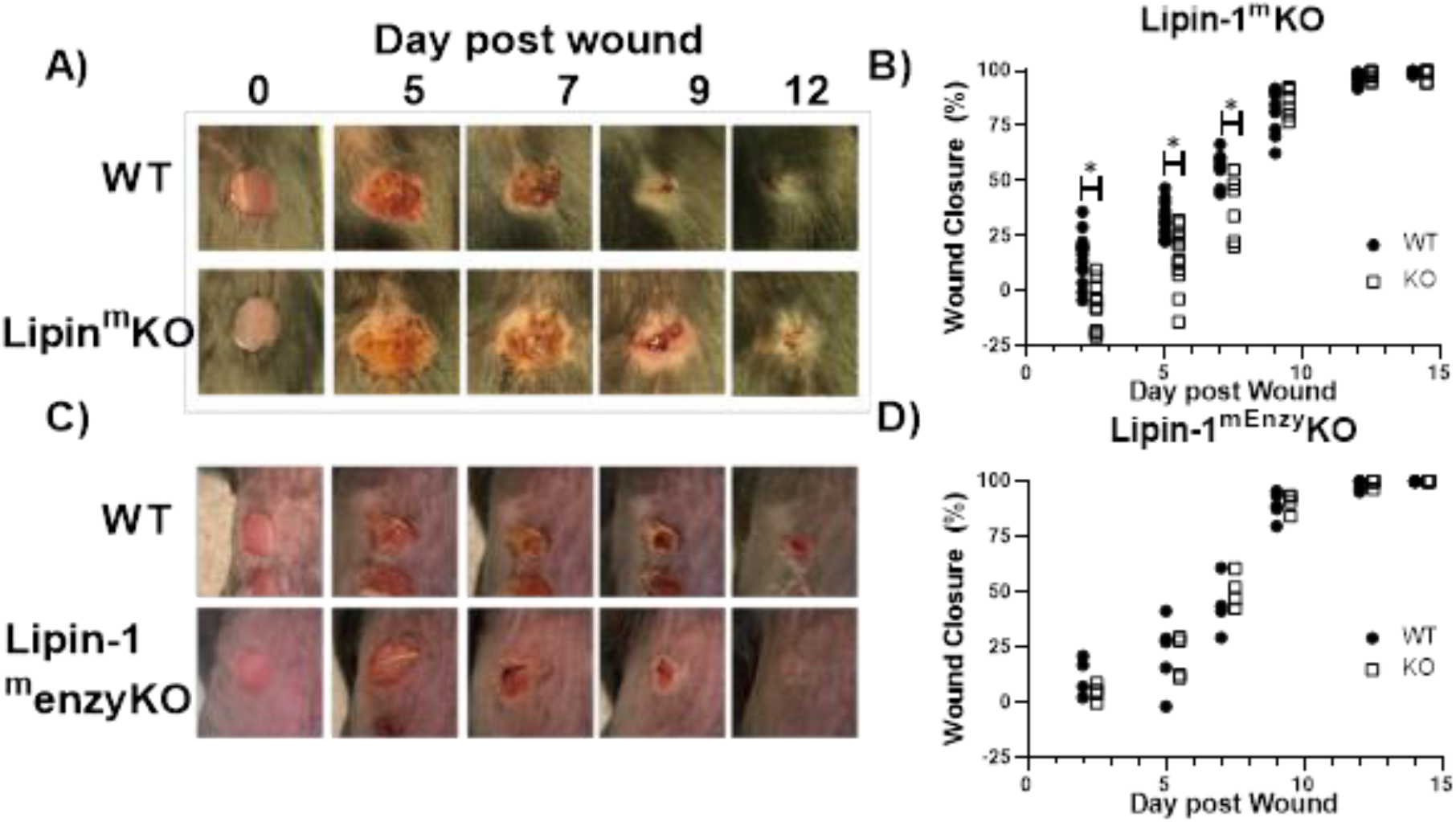
Loss of full lipin-1 delays wound closure. A & C) Representative image of gross lesions. B & D) Percent wounds closure as [(area of original wound – area of actual wound)/area of original wound]x100. Wound measurements were made on days 0, 2, 5, 7, 9, 12, and 14 post-wounding. KO mice are shifted by a half day of graph in order to see differences. (n≥4, *=p≤0.05). Each symbol represents and individual mouse. E) Images of H&E staining of skin. F) Pathology score concerning inflammatory infiltrate and fibrin formation.

Impaired healing was prominent in the early stage of wound healing. Hence further histopathological evaluation (Fig 5A) was performed by H&E staining in isolated wounds from lipin-1^m^KO and littermate control mice at 2- and 5- days post wounding. On day 2, slightly interrupted superficial layer with void spaces were seen at the wound site; scattered mononuclear cells and neutrophils were also observed within the superficial layer in control mice. In Lipin-1^m^KO mice, the superficial layer was poorly bridged with large void spaces and more inflammatory cells (Fig 5A). A very thick crest/scab was evident at the wound area which was highly infiltrated with mononuclear cells and neutrophils indicative of hyper inflammatory phase in Lipin-1^m^KO mice. On day 5 epidermal tongue (depicted by yellow arrow heads) extended towards the center of the wound, indicative of wound bridging and healing in control mice. But, in lipin-1^m^KO mice, the crest region was still thick with large number of immune infiltrates and they lacked a definitive epidermal closure and organization, suggestive of impaired healing. Wound closure (interphase between host tissue and wound depicted by red arrow heads) was also improved in the control mice. Scoring of the stained sections (0-3 inflammatory infiltrate and 0-2 crust thickness) by a blinded pathologist showed no significant difference in inflammatory recruitment in lipin-1^m^KO mice (Fig 5B &C). Even though statistically not significant, Lipin-1^m^KO mice exhibited a high pathology score at day 5 and 12, indicative of increased inflammatory condition at the wound area compared to WT mice. These data suggest that myeloid associated lipin-1 transcriptional co-regulatory activity may be anti-inflammatory in function and is required for early wound healing.

**Figure 5.**
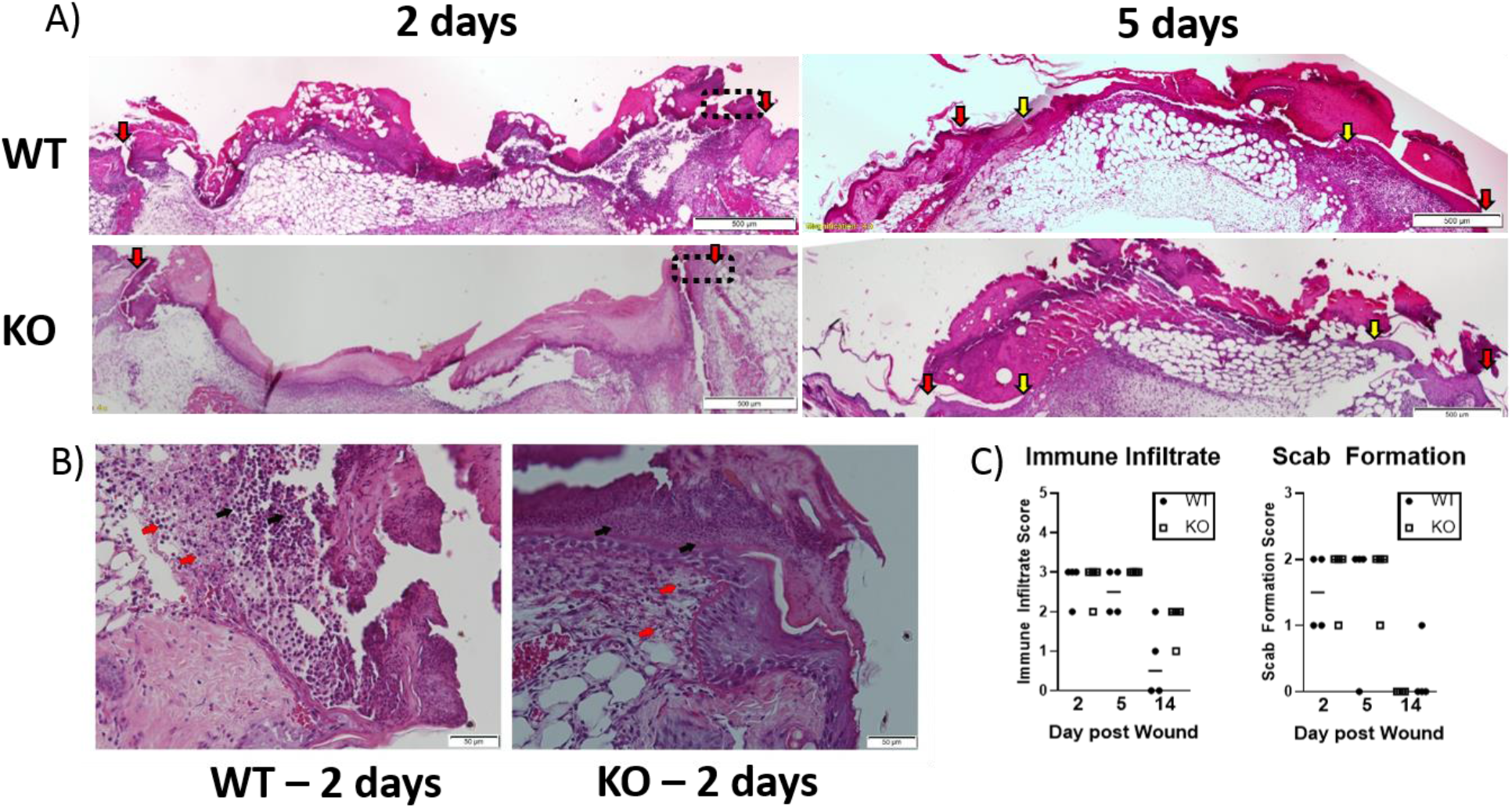
Loss of full lipin-1 delays wound closure. A) H&E depicting epithelial closure of 5mm wounded skin in lipin-1^m^KO and wild type mice post 2 and 5 days of wounding. Red arrow heads indicate host tissue-wound interphase; yellow arrow heads indicate tip of epithelial tongue. B) representative micrographs depicting immune infiltration at the host tissue-wound interphase (highlighted in 5A in rectangular box) post 2 days of wounding. Red arrow points towards monocytes and black arrow points towards neutrophils. C) Pathology score concerning immune cell infiltration and scab thickness (n=4). Scale bar A) = 500μm, B) = 50μm.

### Lipin-1 deletion does not alter myeloid immune composition

Loss of lipin-1 could potentially influence development of myeloid cells or myeloid mediated systemic responses. We examined myeloid population in the spleen and the blood to see if there were any alterations that may explain the delay in wound healing. We isolated the spleen and blood at days 2, 5, and 14 post-wounding. Cells were isolated and stained with a panel of antibodies to quantify macrophages, monocytes, PMNs, and Ly6C^+^ monocytes. We included Ly6C staining as Ly6C^hi^ and Ly6C^lo^ can both contribute to wound healing [26]. We observed no significant difference between lipin-1^m^KO and litter mate control mice in any myeloid cell population analyzed in the blood or spleen (Fig 6A&B). In addition to monitoring cellular responses, we also examined serum cytokine concentration from mice 2 days post-wounding. We chose day 2 as this day correlated with the biggest difference in wound size. No differences were noted in serum cytokine responses between lipin-1^m^KO and littermate control mice (Fig 6C). These data suggest that myeloid associated lipin-1 activity that contributes to wound healing is likely mediated in the local environment rather than systemically.

**Figure 6.**
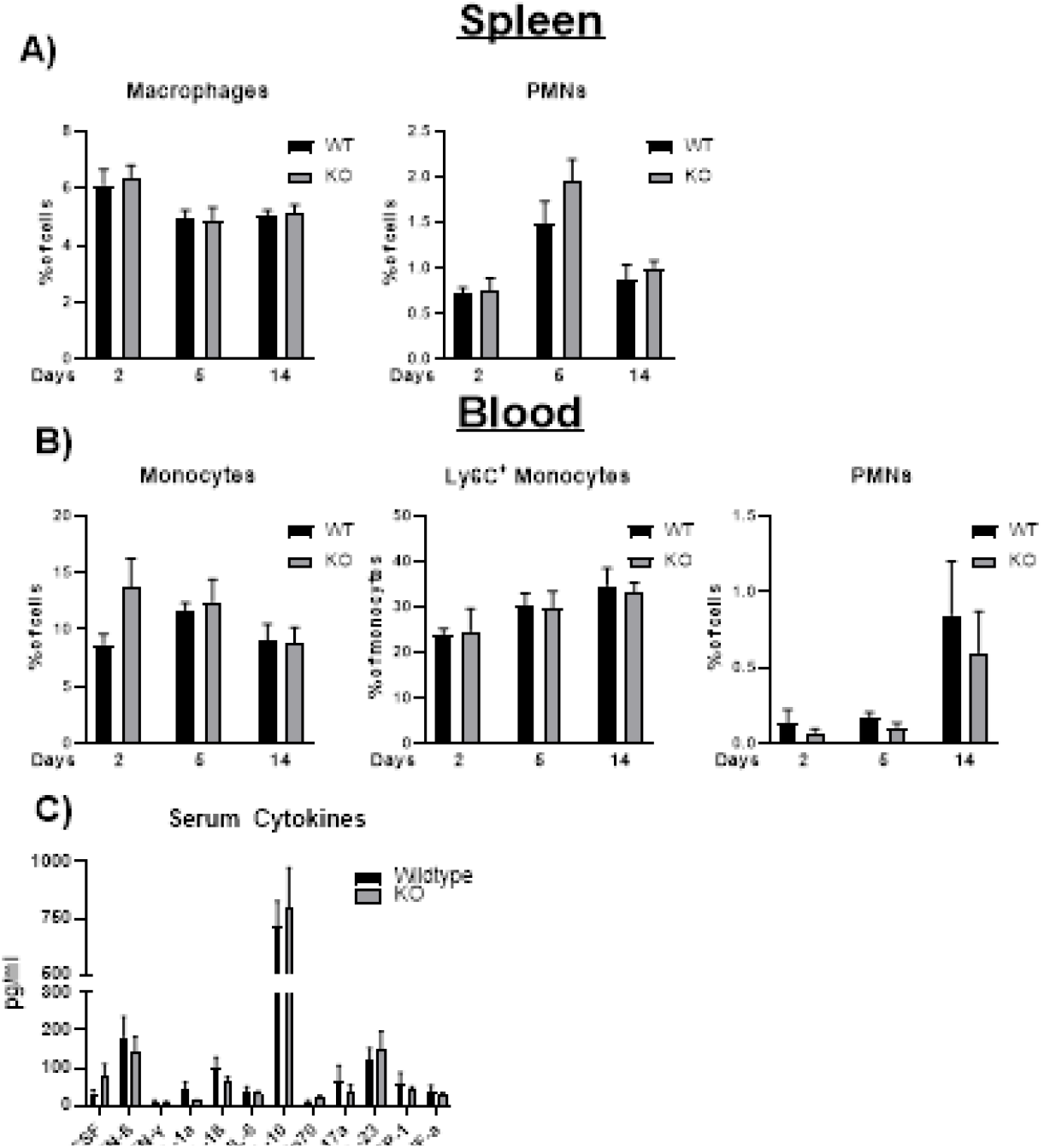
Loss of lipin-1 does not alter systemic immune responses. A) Splenocytes and B) blood were stained with a panel of antibodies to quantify monocyte/macrophage and PMN populations. Myeloid populations were defined as CD45^+^, CD3^−^CD19^−^ CD11b^+^. PMNs were CD11b^+^ Ly6g^+^ and monocytes were CD11b^+^Ly6G^−^. n≥12. C) Cytometric bead arrays to quantify cytokine n≥3 concentrations in serum taken from mice 2 days after wounding n≥4.

## DISCUSSION

Using our lipin-1^m^KO and lipin-1^mEnzy^KO mice, we provide evidence that lipin-1 transcriptional co-regulator activity contributes to wound healing macrophage polarization. Only macrophages lacking the entire lipin-1 protein failed to fully polarize or gain phagocytic capabilities in response to IL-4 which are both activities critical to wound healing function [5]. Impaired healing of full excision wound was also observed in lipin-1^m^KO mice without any alteration in systemic myeloid immune composition. Combined, these data illuminate the contribution of lipin-1 transcriptional co-regulatory activity to macrophage wound healing function.

IL-4 binding to the IL-4R leads to phosphorylation and activation of STAT6. STAT6 binds to DNA promoters that leads to recruitment of PPARγ:RXR transcription factors to promote gene expression in macrophages [2; 22]. PPARγ is highly expressed during macrophage differentiation and its expression is a prominent feature of IL-4 stimulated macrophages [27]. In adipocytes and hepatocytes, lipin-1 binds to and augments the activity of both PPARα and PPARγ [14; 16; 28]. The removal of exons 3 and 4 of lipin-1 in our lipin-1^mEnzy^KO mice results in truncated lipin-1 that lacks enzymatic activity but retains the ability of lipin-1 to bind to transcription factors such as PPARα and PPARγ [16]. BMDMs from the lipin-1^mEnzy^KO mice had equivalent expression of IL-4 elicited genes as WT BMDMs suggesting that lipin-1 enzymatic activity is dispensable for IL-4 mediated gene expression. Removal of exon 7 from lipin-1 in our lipin-1^m^KO mice causes a missense protein leading to loss of lipin-1 and both activities [17]. The lipin-1^m^KO BMDMs did not have a defect in either IL-4R expression nor IL-4 mediated STAT6 phosphorylation. We did observe a reduction in IL-4 mediated gene expression in lipin-1^m^KO BMDMs. IL-4 mediated signaling and PPARγ activity in macrophages can enhance phagocytosis [23; 29]. We observed a reduction in phagocytosis in IL-4 stimulated lipin-1^m^KO BMDMs. Taken together these data suggest to us the lipin-1 transcriptional co-regulator activity is required for IL-4 mediated gene expression and is likely via lipin-1 regulation of PPARγ activity in macrophages. Future work will be to better demonstrate lipin-1 mediated regulation of PPARγ activity in macrophages.

Macrophage polarization is critical for effective *in vivo* wound healing where the number and phenotype of the resident and recruited macrophages determine the extent of healing [30]. Up to 1 day after wounding, pro-inflammatory macrophages initiate an acute inflammatory response and during late phase, wound healing macrophages promote angiogenesis and tissue formation [31]. The loss of lipin-1 is also known to impact pro-inflammatory macrophage polarization via its enzymatic activity [7; 9]. Mice lacking lipin-1 enzymatic activity from their myeloid cells have reduced pro-inflammatory responses. We observed no defect in wound closure in lipin-1^mEnzy^KO mice compared to litter mate controls. In contrast, mice lacking both lipin-1 activities had a defect in wound closure, suggesting that myeloid specific lipin-1 transcriptional co-regulator activity is critical for wound closure. Taken together with our *in vitro* data we would propose this is due to failure in wound healing polarization.

PPARγ KO mice exhibit a significant delay in wound healing at early healing time points (days 5 and 7) due to compromised granulation, collagen deposition, and angiogenesis [5]. Delayed dorsal skin wound healing was exhibited by lipin-1-mFull KO mice at day 2, ,5 and 7 post injury. Results from our lipin-1^m^KO mice are similar to PPARγ KO mice. We would suggest that defective wound healing in lipin-1^m^KO mice may be due to alteration in PPARγ activity, as supported by our *in vitro* gene expression data. During the later stage of healing at day 14, the lipin-1^m^KO exhibited comparable healing to that of WT mice. A similar healing observed, where overall wound closure is the same but early healing is defective has been reported in fibrinogen-deficient mice [32]. Fibrinogen is a component of the extracellular matrix promoting wound closure. IL-4 stimulated macrophages regulate extracellular matrix remodeling through the induction of extracellular matrix production and clearance of extracellular matrix via phagocytosis [24]. These data suggest to us that lipin-1 transcriptional co-regulatory activity likely promotes proper macrophage function and contributed to wound closure.

We propose that the loss of wound healing observed in our lipin-1^m^KO mice is due to loss of transcriptional co-regulatory activity from monocytes and macrophages. However, *LysM*-Cre was used to knockout lipin-1 in our mice, and *LysM* expression is not restricted to monocytes and macrophages. Analysis of *LysM*-Cre–mediated gene deletion demonstrates gene excision in dendritic cells (DC) and granulocytes, as well as monocytes and macrophages [33; 34].We suggest that loss of lipin-1 in DC is not responsible for the difference in wound closure in our lipin-1^m^KO mice. DCs enhance T cell/B cell responses, rather than innate immune responses, and our difference in wound closure is more prevalent in earlier phase of healing (likely prior to T cell responses). The contribution of lipin-1 to DC function is completely unknown and needs to be looked at in the future. Neutrophils clearly contribute to wound healing [35]. However, lipin-1 is not readily detected in neutrophils, suggesting that lipin-1 has little contribution to neutrophil function [7]. Thus, we propose that the loss of lipin-1 transcriptional co-regulatory activity in monocytes and macrophages is most likely responsible for the reduction in wound closure.

Our data highlights the role of lipin-1 transcriptional co-regulator activity within macrophage function, specifically for wound healing polarization. Furthermore, we provide evidence that the lack of myeloid-associated lipin-1 transcriptional co-regulator activity has *in vivo* consequences. Macrophage responses are now recognized to play crucial roles in a diverse array of pathologies like atherosclerosis, arthritis, osteoporosis and sterile inflammation. Future work will be needed to better understand the mechanisms by which lipin-1 transcriptional co-regulatory activity drives macrophage function in different pathological conditions.

## Data Availability Statement

The raw data supporting the conclusions of this manuscript will be made available by the authors, without undue reservation, to any qualified researcher.

## Ethics Statement

All animal studies were approved by the LSU Health Sciences Center-Shreveport institutional animal care and use committee. All animals were cared for according to the National Institute of Health guidelines for the care and use of laboratory animals.

## Author Contributions

SC performed experimental work, data analysis, and wrote the manuscript. RMS, CMRB, AY RM and RSS assisted with experimental design, experimental work and data analysis. CMRB assisted with experimental work and data analysis. RC and BNF provided critical reagents necessary to complete and finalize experiments. MDW conceived the idea, designed the study, obtained fund, analyzed and interpreted data, and wrote and revised the manuscript. All authors were involved in the final approval of the manuscript.

## Acknowledgments

We would like to thank David Custis for his help with running samples related to flow cytometric experiments, Deshaun Blankenship for sectioning all tissues used in this study and Gabrielle Gahn for determining genotypes of all mice.

